# MtrA regulation of essential peptidoglycan cleavage in *Mycobacterium tuberculosis* during infection

**DOI:** 10.1101/2021.02.25.432019

**Authors:** Eliza J. R. Peterson, Aaron N Brooks, David J. Reiss, Amardeep Kaur, Wei-Ju Wu, Robert Morrison, Vivek Srinivas, Serdar Turkarslan, Min Pan, Warren Carter, Mario L. Arrieta-Ortiz, Rene A. Ruiz, Apoorva Bhatt, Nitin S. Baliga

## Abstract

The success of *Mycobacterium tuberculosis* (Mtb) is largely due to its ability to withstand multiple stresses encountered in the host. Here, we present a data-driven model that captures the dynamic interplay of environmental cues and genome-encoded regulatory programs in Mtb. The model captures the genome-wide distribution of *cis*-acting gene regulatory elements and the conditional influences of transcription factors at those elements to elicit environment-specific responses. Analysis of transcriptional responses that may be essential for Mtb to survive acidic stress within the maturing macrophage, identified regulatory control by the MtrAB two-component signal system. Using genome-wide transcriptomics as well as imaging studies, we have characterized the MtrAB circuit by tunable CRISPRi knockdown in both Mtb and the non-pathogenic organism, *M. smegmatis* (Msm). These experiments validated the essentiality of MtrA in Mtb, but not Msm. We identified that MtrA regulates multiple enzymes that cleave cell wall peptidoglycan and is required for efficient cell division. Moreover, our results suggest that peptidoglycan cleavage, regulated by MtrA, is necessary for Mtb to survive intracellular stress. Further, we present MtrA as an attractive drug target, as even weak repression of *mtrA* results in loss of Mtb viability and completely clears the bacteria with low-dose isoniazid or rifampicin treatment.

## INTRODUCTION

*Mycobacterium tuberculosis* (Mtb) infection occurs by inhalation of bacilli-containing aerosols. Mtb initially colonize the lungs, where alveolar macrophages phagocytize the bacteria and sequester them in a phagosome. As Mtb navigate their intraphagosomal environment, they rely on gene regulatory networks (GRNs) that have evolved to evade the bactericidal properties of the maturing phagosome (e.g., acidic pH, reactive oxygen and nitrogen species, nutrient deprivation). Building accurate and comprehensive GRN models can inform how Mtb survives the myriad of stresses within the host environment. A fully characterized GRN of Mtb would also enable actionable hypotheses for identifying novel therapeutic strategies that could disrupt Mtb’s successful tolerance to stress and better control infection.

The GRN structure is encoded in an organism’s genome as transcription factor (TF) binding sites, referred to as gene regulatory elements (GREs). GREs, ∼6-20 nucleotide DNA sequences, are invariant to environmental stresses. However, environmental and genetic perturbations alter the affinity of TFs to bind GREs, which in turn modulate the transcriptome in a condition-specific manner. Therefore, the goal of reconstructing a GRN is to produce an unbiased genome-wide map of GREs, including information about what regulators bind to those sequences, in what contexts they are bound, and, importantly, how TF-binding throughout the genome ultimately influences cellular physiology.

We previously reported models of *Escherichia coli* and *Halobacterium salinarum*^1^ that realized these goals of GRN inference. The models were constructed with EGRIN 2.0, which provided a number of advancements to the previous Environment and Gene Regulatory Influence Networks (EGRIN)^2^. EGRIN models, learned by *cMonkey*^3,4^ and *Inferelator*^5^, have accurately predicted conditional regulatory interactions in a number of organisms including Mtb^6-13^; yet, these network models had some limitations in probabilistically uncovering conditional co-regulation of genes by specific TFs and GREs, especially in under-represented environmental conditions in the transcriptome compendium. Therefore, we developed EGRIN 2.0 to overcome these shortcomings and model the organization of GREs within every promoter and link the contexts in which they act to the conditional co-regulation of genes. Importantly, the improvements and algorithmic components of EGRIN 2.0 have been rigorously tested and validated in prior studies^1,3-5^; readers are encouraged to refer to the original papers for more detail. As such, this study was able to focus on revealing regulatory behaviors that are critical to Mtb pathogenesis, with experimental validation of model-based predictions.

One of the earliest environmental cues encountered by Mtb during infection is a drop in pH during maturation of the macrophage phagosome. In fact, Mtb responds transcriptionally to the acidic pH of the macrophage within 20 minutes of phagocytosis^14^, manifesting in a number of physiological responses, including slow growth (∼pH 6.5) or complete growth arrest (∼pH 5.0)^15,16^. The growth arrested state of Mtb is accompanied by complex changes (e.g., cell wall structure, metabolic activity, target abundance) that can affect the pathogen’s susceptibility to antibiotics.

Here, we have uncovered the regulatory programs that control Mtb’s transcriptional response to diverse host-relevant environmental cues, to understand what makes Mtb such a successful and difficult to treat pathogen. We constructed an EGRIN 2.0 model for Mtb and used the mechanistic detail captured to discover regulatory mechanisms responsible for the pathogen’s pH-adapted lifestyle. Specifically, using EGRIN 2.0 we formulated the hypotheses that the response regulator, MtrA conditionally modulates cell division during acidic stress by regulating multiple cell wall peptidoglycan (PG) cleavage enzymes. Using tunable CRISPRi gene silencing to knockdown *mtrA*, we have validated the EGRIN 2.0-predicted regulatory activity of MtrA, and demonstrate that even minimal *mtrA* knockdown is lethal to Mtb and results in complete clearance with treatment of frontline antibiotics. Together, these results describe the extreme vulnerability of the MtrA regulon and its potential as a novel therapeutic target.

## RESULTS

### Constructing EGRIN 2.0 for Mtb

EGRIN 2.0 is an ensemble framework that aggregates associations across genes, GREs, and environments from many individual EGRIN models, each trained on a subset of gene expression data. The aggregated, post-processed ensemble of EGRIN models is referred to as EGRIN 2.0, which details the organization of GREs within every promoter and identifies conditionally <u>co</u>-<u>re</u>gulated gene <u>m</u>odules, known as corems. The Mtb EGRIN 2.0 model was constructed using the same workflow as the previous EGIRN 2.0 models (**Figure 1A**), with a few advancements: (1) network inference was done with the *cMonkey2* biclustering alogirthm^4^; (2) use of TF-target gene interactions with physical binding (ChIP-seq data)^17^ to inform the set-enrichment scoring module in *cMonkey2*; and (3) manually defined condition blocks to subset the gene expression data. The compendium of gene expression data used for training Mtb EGRIN 2.0 was collected from publicly available datasets and previously used to construct the Mtb EGRIN model^7^. Within the expression compendium, transcriptome profiles of Mtb cultured in conditions of osmotic stress, low iron, carbon monoxide (CO) stress, rifampicin treatment, exposure to lung surfactant or low pH were represented in <2% of the 1900 transcriptomes in the compendium (**Figure 1B**). However, these condition blocks and all condition blocks were each included in ∼20% of *cMonkey2* runs aggregated into EGRIN 2.0 (**Figure 1B**). Thus, the ensemble-based method of EGRIN 2.0 can effectively up-weight rare conditions and reveal context-dependent regulatory interactions that occur infrequently in the data. We compared the detection of enriched gene sets (**Figure 1C**) from these rare conditions within biclusters inferred by Mtb EGRIN and corems identified by Mtb EGRIN 2.0. With the exception of CO stress, corems were significantly better at detecting the enriched gene sets from rare conditions **(Figure 1D**). It is possible that associations related to CO stress would have been better identified if the condition block was consistently grouped with low oxygen (O_2_) and nitric oxide (NO) stress transcriptomes, as CO, O_2_, and NO are sensed concurrently during Mtb infection^18^. Overall, Mtb EGRIN 2.0 better captures conditional co-regulation of genes from rare environments. Given that low pH conditions represent only a small portion of the gene expression data, constructing an EGRIN 2.0 model for Mtb was imperative for this study (**Figure 1E**).

**Figure 1.**
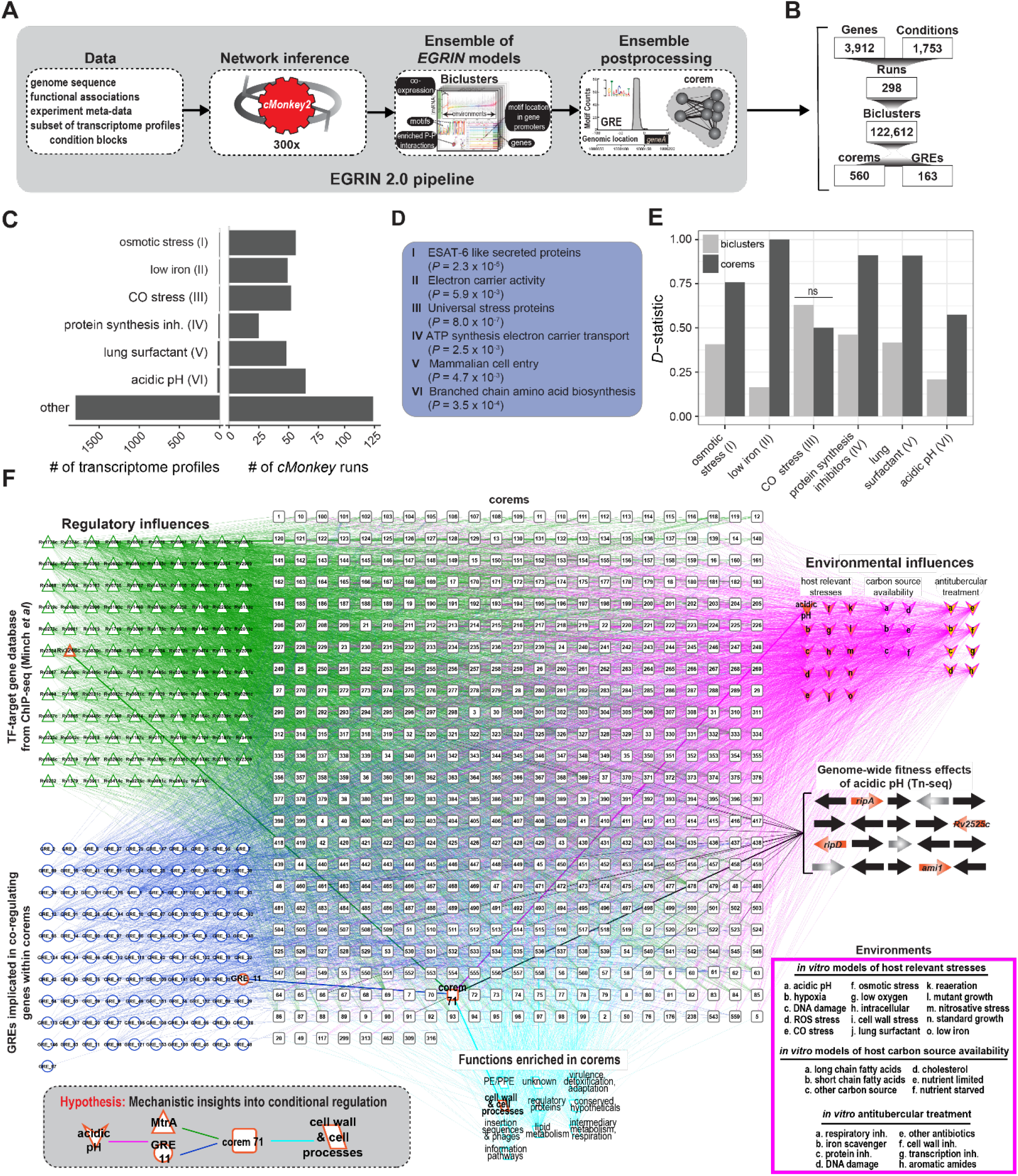
Overview of EGRIN 2.0 model for *M. tuberculosis*. (**A**) The *cMonkey* algorithm was applied many times to subsets of gene expression data from a large compendia of transcriptome profiles to construct many individual EGRIN models. The cMonkey biclustering algorithm identified sets of genes that have co-expression under a subset of experimental conditions, have a common motif in their promoters and are enriched in protein–protein (P–P) interactions. Individual EGRIN models were integrated into an ensemble and mined to construct the final EGRIN 2.0 model, defining overlapping co-regulated sets of genes (corems) that are statistically associated with specific gene regulatory elements (GREs). (**B**) A summary of the counts for each feature in the EGRIN 2.0 model for Mtb. (**C**) Condition blocks representing < 2% of all transcriptome profiles and their inclusion in cMonkey runs. (**D**) General theme of most significant functional cluster defined by DAVID^73^ for each condition block shown in (C). (**E**) Enrichment of functional clusters in (D) for biclusters (EGRIN models containing relevant condition block) and corems of EGRIN 2.0 model, using one-tailed Komogorov-Smirnov (KS) test. We report the KS *D*-statistic. ns, not significant. (**F**) The EGRIN 2.0 model of Mtb with 560 corems, each of which is statistically associated with GREs (blue circles) or transcription factors (TFs) from ChIP study by Minch *et al*^*17*^ (green triangles). Corems are also statistically associated with environmental influences (pink diamonds) and general mycobacterial functions (teal parallelograms) defined by MycoBrowser^74^. We highlight a signal path through the network that is discussed in detail (bold edges and nodes): genes essential for growth at acidic pH are significantly enriched in corem 71, which is predicted to be regulated by MtrA and GRE #11 and implicated in processes related to the cell wall. The inset illustrates ways in which the network model can be used to make actionable predictions. The diagram was generated with Biotapestry^75^. CO: carbon monoxide. inh: inhibitor(s).

### Genome-wide fitness screening identifies corems required for *M. tuberculosis* adaptation to acidic pH

To identify the regulatory networks involved in Mtb adaptation to acidic pH, we utilized transposon mutagenesis coupled with next-generation sequencing (Tn-seq)^19^. Transposon mutant libraries containing ∼100K individual transposon mutants were generated in Mtb and cultured in either pH 7.0 or pH 5.6 media for 3 days, at which point the surviving bacteria were plated. Transposon gene junctions were amplified and sequenced from the recovered bacteria and *TRANSIT* Tn-seq analysis tool^20^ was used to identify transposon insertion sites and calculate their representation in each condition. We developed new software to quantify, display, and summarize the comparative fitness of genes at acidic pH versus neutral pH (see Methods, **Dataset S1**). The corem fitness at acidic pH was then calculated by averaging the conditional change in fitness scores for the genes in each corem (**Dataset S2**). This analysis revealed corem 71 as being significantly enriched with genes having decreased fitness at acidic pH (permuted Benjamini-Hochberg, BH, corrected *p*-value < 0.001). We performed another Tn-seq fitness screen in the presence and absence of the cell wall-perturbing detergent sodium dodecyl sulfate (SDS). There was no significant decrease in average fitness for corem 71 genes in the presence of SDS (**Figure S1, Dataset S1** and **S2**), suggesting that corem 71 is enriched in genes with reduced fitness specific to acidic pH.

### MtrA regulates corem 71 genes that are required for *M. tuberculosis* adaptation to acidic pH

To find evidence of corem 71 regulation by specific TFs, we investigated the EGRIN 2.0-predicted organization of GREs in the promoters of corem 71 genes. EGRIN 2.0 predicts the frequency of GRE alignment to each genomic position, thus predicting the organization of GREs in gene promoters at nucleotide resolution. EGRIN 2.0 predicts that five GREs are significantly enriched in the promoters of corem 71 genes, with GRE #11 being the most significant (BH corrected *p*-value = 8.8 × 10^−11^) and found in the promoter of all corem 71 genes with reduced fitness at acidic pH (**Figure 2A**). We compared the motif of GRE #11 to motifs that were deciphered through analysis of ChIP-seq mapped TF binding locations^17^. GRE #11 significantly matched (*p*-value < 0.01) the sequence motif of MtrA (Rv3246c) determined by ChIP-seq^17,21^. Moreover, MtrA ChIP-seq binding sites were identified in the promoters of corem 71 genes, including the genes with reduced fitness at low pH, with statistical significance (BH corrected *p*-value < 0.0001). In addition to the genes with reduced fitness at low pH, corem 71 contains 46 genes including the TF, MtrA (**Figure 2B**). MtrA is the response regulator of a two-component system (TCS) with MtrB. MtrAB is essential for *in vitro* growth of Mtb^22,23^, and predicted to be essential for Mtb survival in macrophages^24^ and Mtb growth in mouse spleen^25^ as determined by Tn-mutagenesis based investigations. Overexpression of MtrA has no effect *in vitro* but results in decreased growth in macrophages^26^, although the mechanism is unknown. Similar to other essential TFs, there is little characterization of MtrA’s regulatory function in Mtb. To better characterize MtrA and its potential role in regulation of corem 71 genes and in acidic pH, we used recent advances in mycobacterial CRISPR-interference (CRISPRi)^27^ to knockdown the expression of *mtrA*. We designed five small guide RNAs (sgRNAs) targeting *mtrA* in Mtb that achieved varying levels of gene knockdown with the CRISPR1 Cas9 from *Streptococcus thermpohilus* (**Table S1**)^27^. Two of the Mtb CRISPRi strains (with sgRNA2 and sgRNA3) reached high-level mRNA repression upon the addition of 100 ng/mL anhydrotetracycline (ATc) and resulted in 3-log viability reduction of Mtb (**Figure S2A**), compared to controls plated without ATc. With ATc induction, we also observed very limited growth and severe cell aggregation in broth culture (**Figure S2B**). The other three CRISPRi strains (with sgRNA4, sgRNA5, sgRNA6) achieved lesser degree of *mtrA* knockdown and viability reduction upon ATc addition (**Figure S3A**). These CRISPRi strains, with partial knockdown of *mtrA*, were able to grow in broth culture in the presence of ATc, although still with distinct growth inhibition compared to no ATc controls (**Figure S3B**). The panel of sgRNAs allowed us to tune *mtrA* knockdown and obtain strains with a range of growth properties for further characterization studies. Using sgRNA2 and sgRNA3, which produced high-level knockdown of *mtrA*, we collected samples four days after ATc addition for gene expression profiling by RNA sequencing (RNA-seq). We performed differential expression analysis with respect to uninduced (-ATc) controls and confirmed that CRISPRi knockdown of *mtrA* repressed the expression of many corem 71 genes (**Figure 2C, Dataset S3**). In particular, all of the corem 71 genes with reduced fitness at acidic pH were significantly repressed (*p*-value < 0.01) upon *mtrA* knockdown (**Table 1**). Interestingly, the isoniazid induced genes (*iniBAC*) were significantly up-regulated when *mtrA* expression was silenced (**Figure 2C**). The *iniBAC* genes are induced by a broad range of cell wall biosynthesis inhibitors^28^ and points toward defects in the cell wall when *mtrA* is silenced.

**Table 1.**
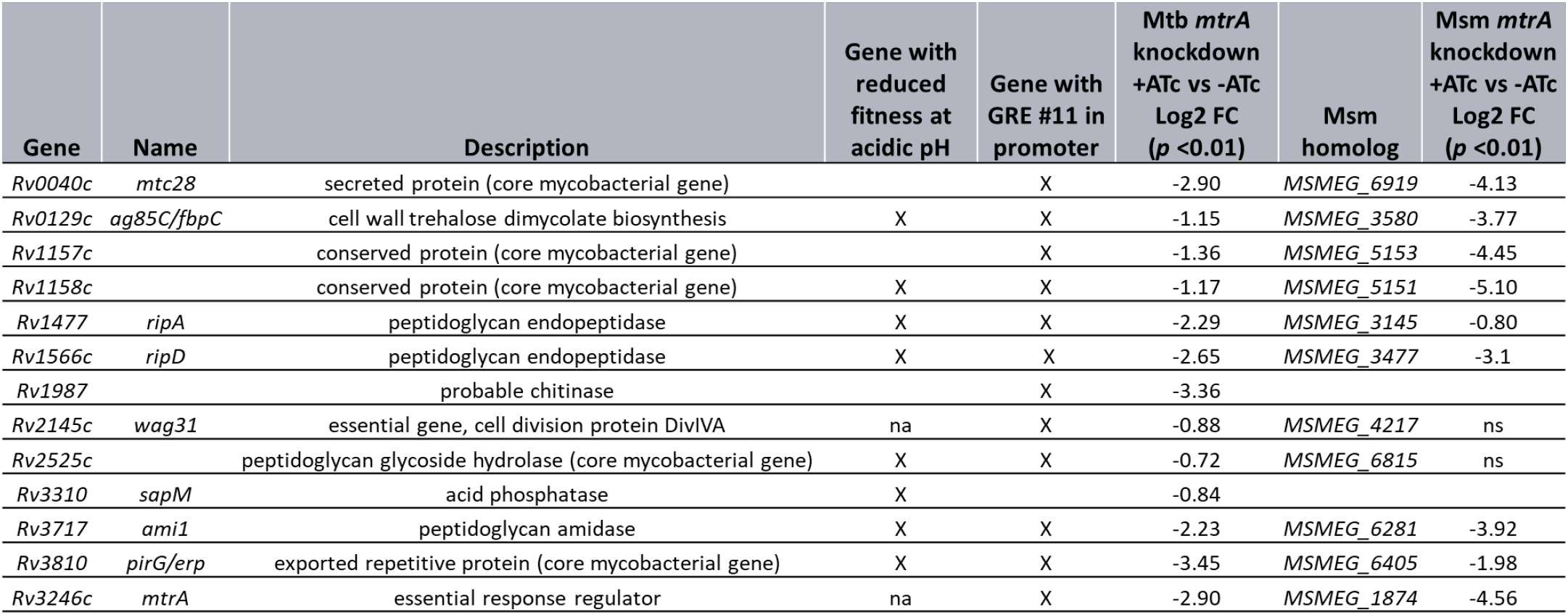
Corem 71 genes with predicted regulation by MtrA. na: not applicable; ns: not significant

**Figure 2.**
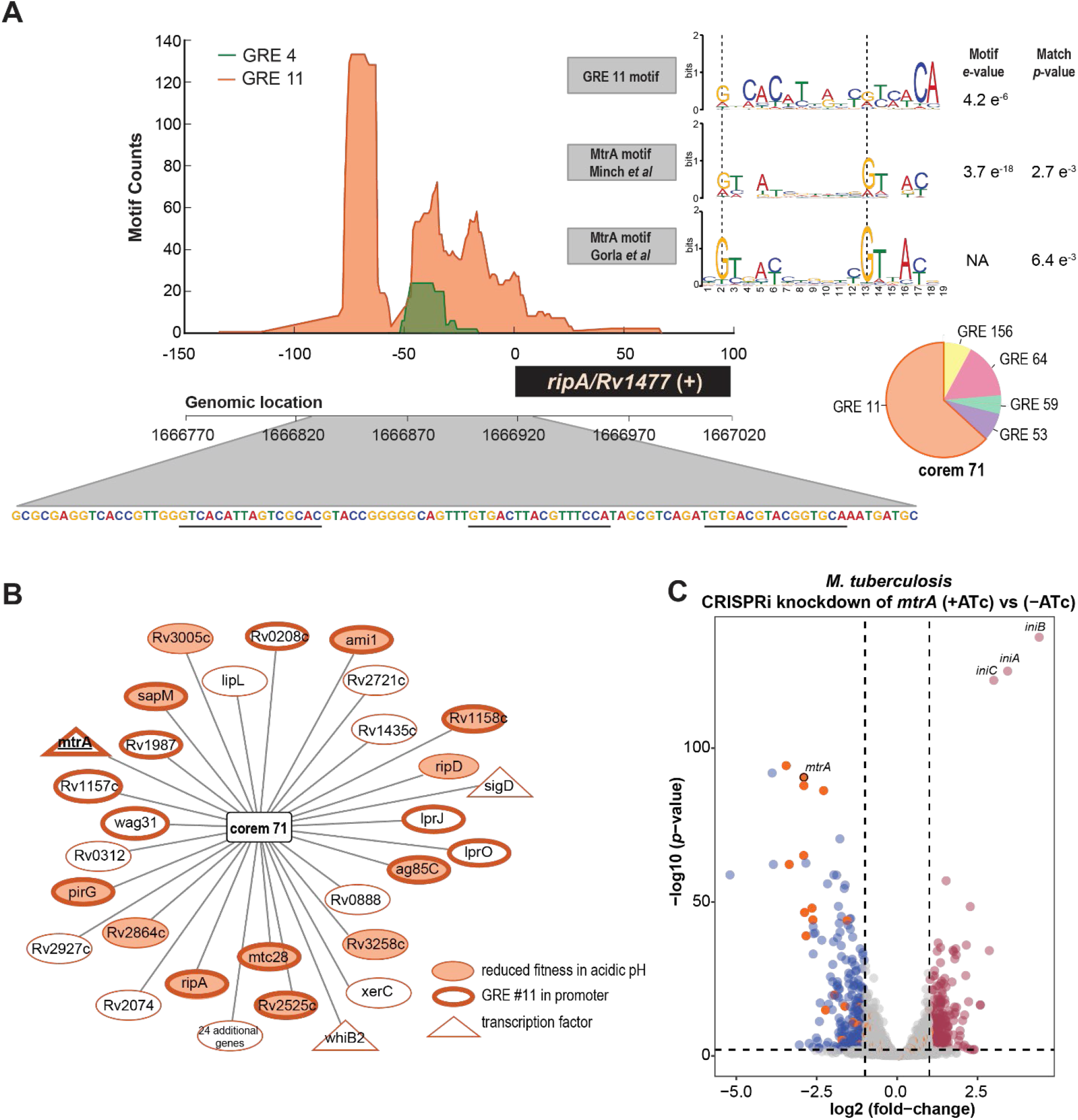
Validation of EGRIN 2.0 predicted regulation of corem 71 genes. (A) Promoter architecture of the *ripA* (Rv1477) promoter predicted by EGRIN 2.0 model. (Top-left) Frequency of GRE alignment to each position in the *ripA* promoter. GREs are indicated by lines, with GRE #11 shaded in orange. (Bottom-left) Genome sequence marked with putative GRE #11 locations. (Top-right) The motif logo of GRE #11 compared with MtrA motifs that were deciphered through analysis of ChIP-seq mapped binding locations. The *e*-values of the motifs and *p*-values from alignment to GRE # 11 carried out with Tomtom^76^ are shown. NA: not available (Bottom-right) Pie chart represents average predicted influence of GREs on the regulation of genes in corem 71. (**B**) Network visualization of genes (ellipses) in corem 71. (**C**) Volcano plot of differentially expressed genes for induced and uninduced CRISPRi knockdown of *mtrA* with sgRNA2 in Mtb. The significantly differentially expressed genes were selected by *p*-value < 0.01 and absolute log2 fold-change > 1. Dots represent different genes, with labels for particular genes of interest. Grey dots are genes without significant different expression, red dots are significantly up-regulated genes (*N* = 266 genes) and blue dots are significantly down-regulated genes (*N* = 240 genes). The orange dots are all genes of corem 71.

Given the sequence similarity of MtrA between *M. smegmatis* (Msm), a non-pathogenic fast-growing mycobacterium, and Mtb, we sought to also characterize *mtrA* knockdown in Msm. We generated CRISPRi strains that achieved equally strong repression of *mtrA* in Msm upon ATc addition. This repression of *mtrA* resulted in a moderate (but significant) growth inhibition of Msm (**Figure S4**), supporting a recent study that found *mtrA* is not essential in Msm, in contrast with the Mtb ortholog^21^. We performed RNA-seq and analyzed differential expression of all genes due to ATc-induced repression of mtrA in the Msm CRISPRi strain (**Figure S5, Dataset S4**). Similar to Mtb, the knockdown of *mtrA* in Msm also resulted in significant repression of the orthologous genes with reduced fitness at low pH (**Table 1**), with the exception of the Rv2525c homolog (MSMEG_6815). Of note in the Msm *mtrA* knockdown strain, was significantly high expression of genes encoding the peptidoglycan (PG) transglycosylases known as resuscitation promoting factors, *rpfB* (MSMEG_5249, log2 fold-change = 2.47, *p*-value < 0.01) and *rpfE* (MSMEG_4643, log2 fold-change = 2.47 and 5.11, respectively, *p*-value < 0.01). RpfB and RpfE are known to interact with Rpf interacting protein, RipA, in Msm, where the protein complex hydrolyzes PG during cell division^29-31^. RipA is found in Corem 71, had reduced fitness in low pH, and was significantly repressed upon *mtrA* knockdown (in both Mtb and Msm). It is possible that increased expression of the *rpfs* (via unknown regulatory mechanism) could sustain PG cleavage in Msm when *ripA* is repressed by MtrA. In contrast, *rpfB* and *rpfE* were not significantly differentially expressed upon *mtrA* knockdown in Mtb. The interaction of RipA and Rpfs, or specifically the lack of interaction in Mtb needs further investigation, and could contribute to the difference in MtrA essentiality between Msm and Mtb. These results suggest that while the regulatory activity of MtrA is largely conserved from Msm to Mtb, subtle differences in the function of this TF that has emerged upon their evolutionary divergence has manifested as a particular vulnerability for the human pathogen.

### MtrA regulon of *M. tuberculosis* is repressed at low pH

To examine the transcriptional response of the MtrA regulon (defined as genes in **Table 1**) at low pH, we grew wild type Mtb to mid-log phase in 7H9 growth medium at neutral pH, washed, then diluted the cells in either neutral or low pH media (7H9-rich media supplemented with 0.05% tyloxapol and adjusted to pH 7.0 or pH 5.6). Following transition to either neutral or acidic media, we monitored growth for 7 days and collected samples at 24 h for transcriptome profiling by RNA-seq. As expected, Mtb growth was stalled at acidic pH (**Figure 3A**). We also observed a significant decrease in MtrA regulon expression at acidic pH compared to neutral pH (**Figure 3B**). This change in expression of MtrA regulon genes, along with the Tn-seq data demonstrating their essentiality at low pH, suggests that reducing but maintaining expression of the regulon genes is important for Mtb’s survival in acidic conditions.

**Figure 3.**
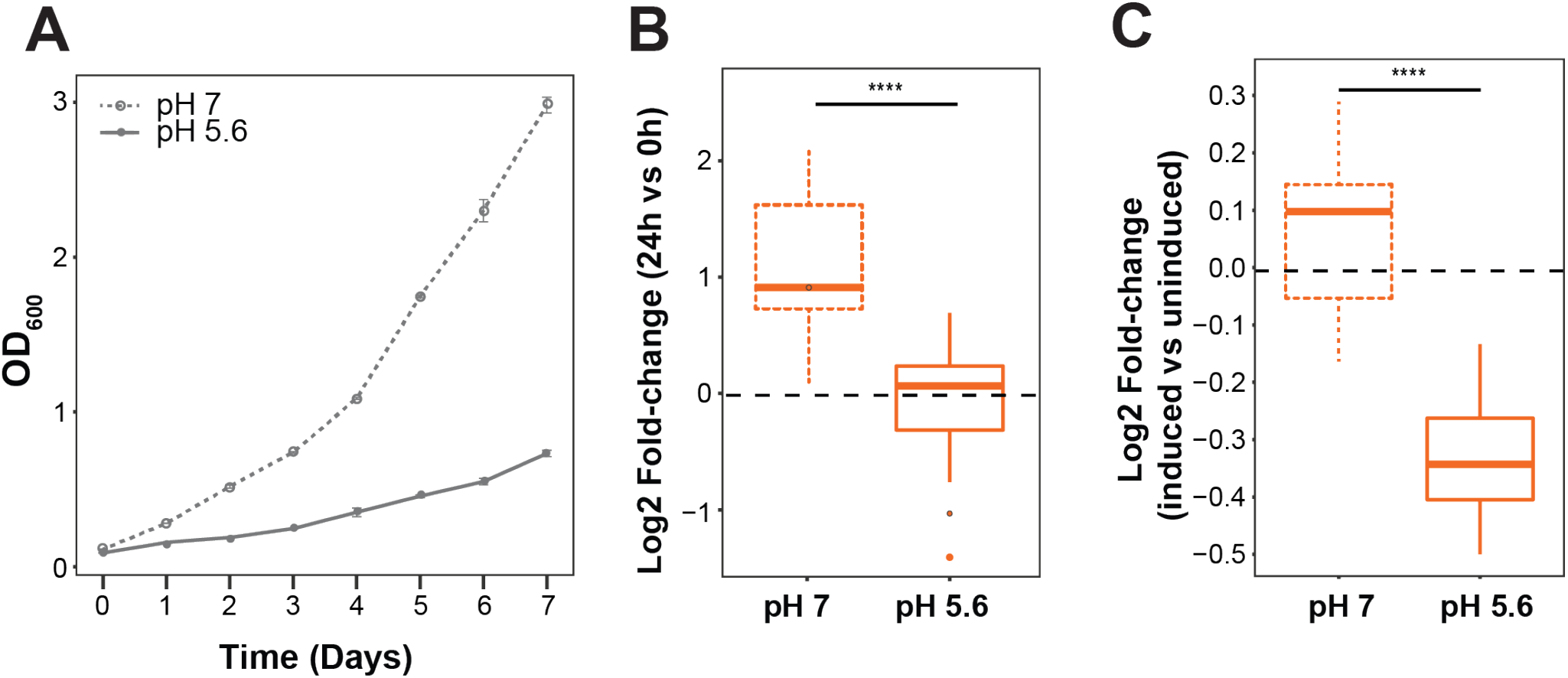
Characterization of corem 71 in acidic pH. (A) Growth of wild-type Mtb H37Rv in 7H9-rich media buffered to either pH 7 (circles, dashed line) or pH 5.6 (squares, solid line). Growth was monitored daily by optical density at 650 nm. Points are the average of three biological replicates and error bars represent standard deviation. (**B**) Boxplot representing log2 fold-change of corem 71 genes with reduced fitness in acidic pH for wild-type Mtb H37Rv in 7H9-rich media buffered to either pH 7 (dotted line) or pH 5.6 (solid line). Data are for three biological samples comparing expression at 24 h vs 0 h. (**C**) Boxplot representing log2 fold-change of Mtb H37Rv with inducible overexpression of *mtrA* for corem 71 genes with reduced fitness. Data are for three biological samples in 7H9-rich media buffered to neutral pH 7 (dotted line) or acidic pH 5.6 (solid line) comparing induced (+ATc) vs uninduced (-ATc) overexpression of *mtrA*. Statistical significance was calculated as *p*-value with Student’s T-test. ****: *p*-value < 0.0001.

Given that the mtrA knockdown strains were growth attenuated and would be further growth inhibited in low pH media, we reasoned that one approach to understand MtrA perturbation at acidic pH would be to overexpress *mtrA*. Using the ATc-inducible overexpression strain previously described^17^, we repeated the transfer of mid-log phase cells to either neutral media or low pH media. After transfer, we induced *mtrA* expression with ATc and collected cells at 18 h for transcriptional profiling by RNA-seq. Addition of ATc resulted in six-fold increase in *mtrA* expression relative to uninduced at neutral pH and four-fold increase in *mtrA* expression at acidic pH (*p*-value < 0.001). While the expression of the MtrA regulon was not affected by overexpression of *mtrA* at neutral pH, it was significantly downregulated at acidic pH (**Figure 3C**). Together, these results suggest that overexpression of *mtrA* during macrophage infection, when Mtb also experiences acid stress, interferes with optimal MtrA levels and possibly its phosphorylation by MtrB^26^, resulting in severe repression of its downstream target genes. These results explain previous observations that *mtrA* overexpression had no effect on *in vitro* growth of Mtb, but abolished its ability to replicate in THP-1 macrophages^26^.

### MtrA knockdown results in *M. tuberculosis* and *M. smegmatis* growth defects

As mentioned above, among the regulatory targets of MtrA (**Table 1**) are four genes with PG hydrolytic activity. These genes, *ripA, ripD, Rv2525c* and *ami1*, encode enzymes that are capable of cleaving PG and contribute to the physical separation of two daughter cells during cell division. RipA is a D,L-endopeptidase, cleaving PG within the peptide stem^32^. RipD also contains an endopeptidase NlpC/p60 domain similar to RipA, but the purified protein demonstrated a non-catalytic PG-binding function^33^. Structural studies suggest a PG glycoside hydrolase function for Rv2525c, although its activity has not been established^34^. Finally, Ami1 contains an amidase_3 domain that cleaves the peptide stem from N-acetyl-muramic acid on the PG glycan backbone^35^. To assess the effect of MtrA regulation on these PG cleavage enzymes and cell division, we used the fluorescent D-alanine analogue HADA, which incorporates into newly synthesized PG. We fluorescently labeled the PG of Mtb *mtrA* knockdown cells containing sgRNA5, which achieved low-level *mtrA* repression, and observed elongated multiseptated cells (**Figure 4A**). Cells with ATc-induced *mtrA* knockdown had an increased median cell length and a larger variation in cell length compared to uninduced cells (**Figure 4B**). Nearly all (∼75%) of the *mtrA* knockdown cells contained at least one septum, while septa were present in only 15% of the uninduced cells (**Figure 4C**). In fact, more than one septum were present in a majority of the *mtrA* knockdown cells, indicating that septa form but the cells fail to divide. Interestingly, the *mtrA* knockdown cells adopted an abnormal curved or bent shape at the poles, which was not found in the uninduced cells (**Figure 4A**). As PG is vital for maintaining cell shape^36^, the curved shape phenotype could be a result of a disrupted PG layer in growing cells. In mycobacteria, growth and division are uncoupled events and the organism continues to grow regardless of the occurrence of cell division^37^. During cell growth, PG cleavage (via PG hydrolases) may be important to allow for incorporation of new PG at the cell poles^38^. It is interesting to speculate that repression of the PG hydrolases in the *mtrA* knockdown strain may affect the integrity of PG in growing cells and lead to the curved shape phenotype. However, further investigation is needed to confirm this hypothesis. Curved, elongated multiseptated cells, with some cells containing branches, were also observed with the Msm *mtrA* knockdown strain (**Figure S6**). While *ripA* (*MSMEG_3145*), *ripD* (*MSMEG_3477*) and *ami1* (*MSMEG_6281*) were significantly repressed in the Msm *mtrA* knockdown strain (**Table 1**), the Rv2525c homolog (MSMEG_6815) was not. This indicates similar, but also unique, roles of MtrA in regulating PG cleavage between Msm and Mtb.

**Figure 4.**
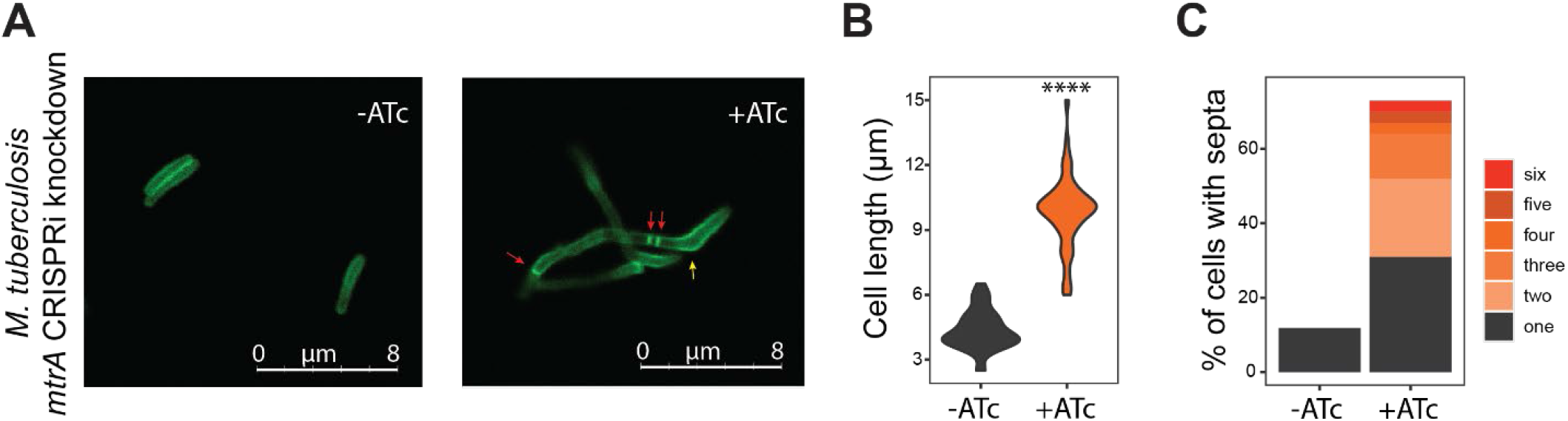
MtrA controls cell division in *M. tuberculosis*. (A) Example micrographs of uninduced (-ATc) and induced (+ATc) CRISPRi knockdown of *mtrA* with sgRNA5 in Mtb. After knockdown, cells were labeled with HCC-amino-D-alanine (HADA) for 20 h. Red arrows indicate multiple septa and yellow arrow indicates the curved shape phenotype. (B) Violin plots showing the cell lengths of uninduced and induced CRIPSRi knockdown of mtrA with sgRNA5 in Mtb. Significance was determined by Student’s T-test. ****: *p*-value < 0.0001. (C) Bar plots representing the percentage of cells that contain 1 or more septa for uninduced and induced CRIPSRi *mtrA* knockdown with sgRNA5 in Mtb. The colored sections within the bars represent the proportions of cells with different numbers of septa. Representative results from two independent experiments are shown.

### MtrA knockdown increases the susceptibility of *M. tuberculosis* to frontline antibiotics

A consequence of defective cell division and growth could be altered susceptibility to antitubercular drugs. We used our Mtb CRISPRi strain with sgRNA5 to determine whether the bactericidal activities of frontline TB drugs are altered when *mtrA* expression is moderately repressed. We compared time-kill curves for the *mtrA* knockdown strain induced with ATc versus uninduced when exposed to sub MIC concentrations of isoniazid and rifampicin. Isoniazid is a frontline antitubercular drug that rapidly kills actively dividing bacilli, whereas rifampicin is a sterilizing drug that kills both replicating and persistent bacilli^39,40^. The two drugs have different mechanisms of action and different chemical properties, yet at concentrations that only partially inhibited growth in the uninduced controls, both drugs were able to completely sterilize the Mtb mtrA knockdown cultures (**Figure 5A**). There were different drug susceptibility patterns observed in Msm *mtrA* knockdown cultures, as measured by disk diffusion (**Figure S7A**). We found that Msm *mtrA* knockdown cultures showed increased susceptibility to rifampicin, carbinicillin and vancomycin. We also tested cycloserine and isoniazid, but saw no difference in susceptibility compared to the control. The results for the Msm mtrA knockdown demonstrated a pattern of increased susceptibility to higher molecular weight antibiotics. Larger molecules may more easily enter the Msm mtrA knockdown cells due to alterations in cell wall composition. To further investigate the changes in drug susceptibility in Msm, we tested the cell wall permeability of the Msm *mtrA* knockdown cells using an ethidium bromide (EtBr) diffusion assay and found *mtrA* repression resulted in reduced EtBr diffusion compared to control (**Figure S7B**). This suggests that the mechanism for increased susceptibility to high molecular weight antibiotics in Msm is not simply due to increases in cell wall permeability. Other drug transport mechanisms such as porin-like proteins may be affected in the Msm *mtrA* knockdown cell.

**Figure 5.**
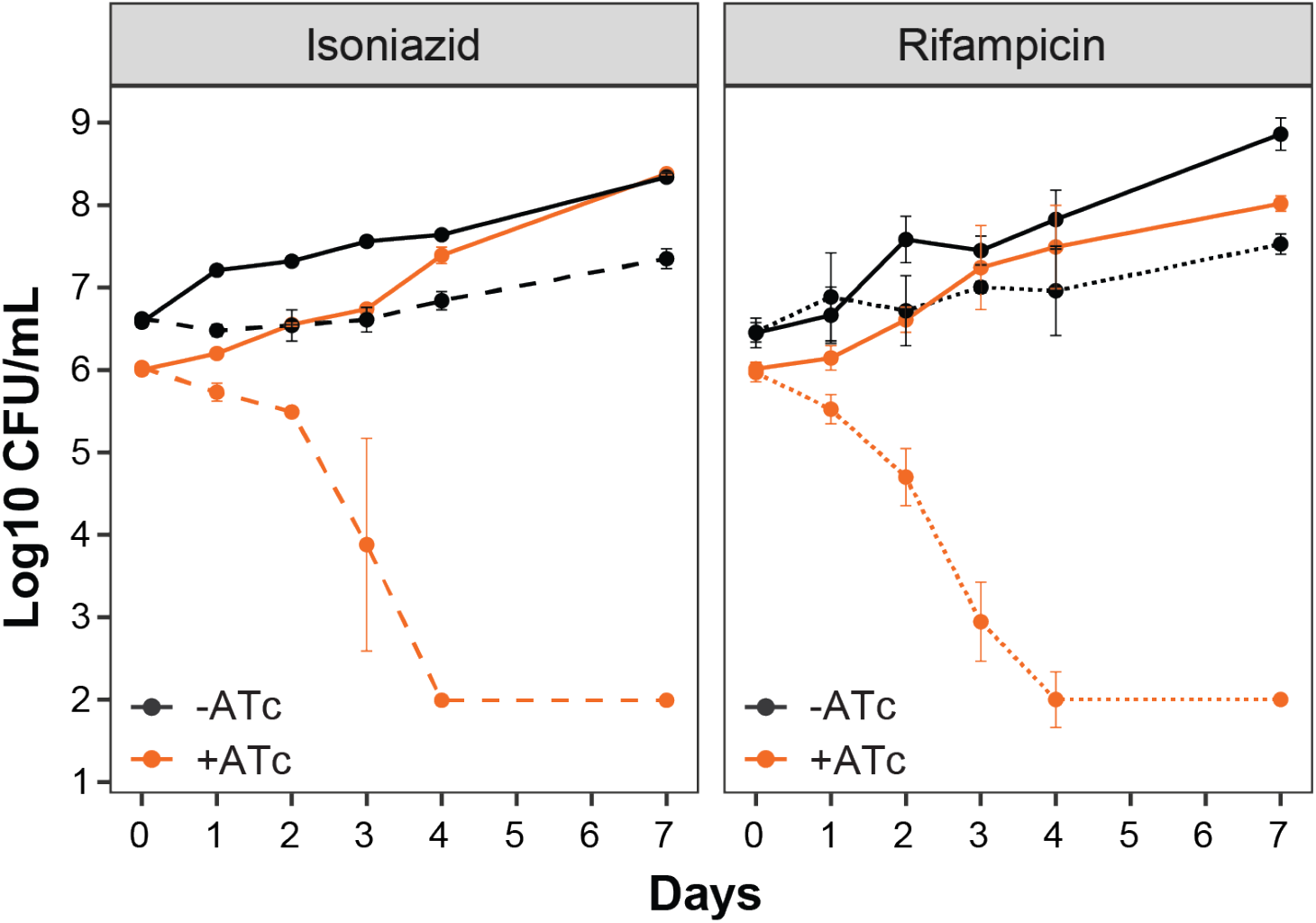
The knockdown of *mtrA* sensitizes *M. tuberculosis* to frontline TB drugs. Colony forming units (CFU) over a period of 7 days for untreated (solid lines) and 0.18 μg/mL isoniazid or 0.006 μg/mL rifampicin treated (dotted lines) liquid cultures of uninduced (black) and induced (orange) CRIPSRi knockdown of mtrA with sgRNA5 in Mtb. Error bars show the standard deviation from three biological samples. Representative results from two experiments are presented.

## DISCUSSION

The EGRIN 2.0 model for Mtb reveals, with unprecedented detail, how the pathogen tailors transcriptional responses to survive stresses experienced within the host. The model captures the genome-wide distribution of GREs within each promoter and statistically significant gene associations in corems from host-relevant conditions. The ensemble network inference methodology allows detection of regulatory events for conditions that represent only a small portion of the entire data set, such as acidic pH, a stress Mtb faces in the maturing macrophage phagosome. We performed extensive experiments to validate a model prediction that genes controlled by the MtrAB regulatory circuit are essential for Mtb at acidic pH. We demonstrated that MtrA regulation of PG cleavage is required for Mtb cell division. Further, we showed that even weak repression of *mtrA* results in significant growth inhibition and completely clears the bacteria with low-dose isoniazid or rifampicin treatment.

Mtb has successfully exploited macrophages as its primary niche *in vivo*, evolving the capability to withstand a myriad of stresses with the macrophage, including acid stress. Several proteins have been implicated in Mtb’s response to acid stress, such as the two-component system PhoPR^15,16,41^, the outer membrane protein OmpATB^42^, and the PG hydrolase RipA^43^. Our work also suggests that PG cleavage by MtrA-regulated PG hydrolases is necessary for Mtb to reside in the acidic phagolysosome. The essentiality of PG cleavage during acid stress points towards residual cell division or PG hydrolysis for the incorporation of new PG, even when Mtb growth is reduced. It has been proposed that replication during stress may favor bet-hedging by distributing damaged macromolecules between progeny cells, thus creating heterogeneous subpopulations with altered fitness and the capacity to survive stress^44,45^. Alternatively, the PG cleavage enzymes may permit PG rearrangement or repair, which could lead to cell wall changes for nutrient uptake^46^ or the maintenance of pH homeostasis^47^ during acid stress.

The maturing macrophage phagosome acquires bactericidal properties, such as acidic pH, to restrict and kill invading pathogens. However, it has been shown that during infection in mice, some macrophage lineages control Mtb better than others^48-50^. Analysis of Mtb’s transcriptional response from alveolar macrophages, which promote Mtb growth, and monocyte derived macrophages, that restrict Mtb growth^51^, finds that the MtrA regulon is significantly down-regulated in monocyte derived macrophages compared to alveolar macrophages (log2 fold-change = -1.1, *p*-value = 6.76 × 10^−5^). This suggests that monocyte derived macrophages contain acidified phagolysosomes that trigger MtrA-mediated repression of PG cleavage. This also supports the notion that monocyte derived macrophages may be innately more capable of restricting bacterial growth compared to alveolar macrophages, since repression of mtrA and target genes would also inhibit cell division. We propose that inhibiting MtrA could restrict Mtb growth in both macrophage populations and promote clearance from the host.

Our study reveals that Mtb cell division, particularly PG cleavage, is transcriptionally regulated by MtrA. We have demonstrated that the MtrA regulon includes multiple PG hydrolases, selected for co-regulation by evolution. Interestingly, corem 71 includes three other PG remodeling enzymes (*dacB1* (Rv3330), penicillin-binding lipoprotein (Rv2864c), and the amidase, *ami3* (Rv3811)), that were not differentially expressed with *mtrA* knockdown in Mtb. This demonstrates how corems group together genes that are functionally related even though their co-regulation is mediated by different mechanisms^1^. Despite their similar enzymatic functions, the PG cleavage enzymes controlled by MtrA could fulfill individually important roles in specific conditions. Recent work established the contextual redundancy of the MtrA regulatory targets, *ripA* and *ami1* in Mtb^52^. The authors showed that Ami1 sustains *in vitro* cell division in cells lacking *ripA*, and vice versa, but that this redundancy is not sufficient for replication and persistence of Mtb in the host^52^. While we demonstrated that MtrA coordinates the transcription of both PG cleavage genes (along with *ripD* and *Rv2525c*), the conditional nature of their redundancy could be related to interacting proteins that may be associated with specific conditional or temporal events during infection. Regardless of their individual roles, their coordinated regulation offers unique opportunity to target multiple PG modifying enzymes by inhibiting MtrA with a drug. In fact, small molecules have been identified that inhibit the DNA binding activity of MtrA^53^. The compounds were found to inhibit the expression of *mtrA* and restrict Mtb replication *in vitro* and within macrophages. As such, there exists chemical space for the development of potent MtrA inhibitors.

Earlier studies proposed a handful of MtrA regulatory targets (*dnaA, ripA, fbpB, ftsI* and *wag31*) using Msm or Mtb strains overproducing wild-type and mutant MtrA^21,26,54,55^. Our model and experimental results confirmed the previously reported MtrA targets, *ripA* and *wag31*. Additionally, several recent studies have evaluated MtrA regulatory targets using global ChIP-seq analysis with Mtb strains overexpressing C-terminal FLAG-MtrA^17^, N-terminal His-tagged MtrA^56^, or proposed phosphorylation competent MtrA protein (replacement of tyrosine with cysteine at position 102)^21^. Among the 14 putative MtrA targets identified across these studies (**Table S2**), we observed significant differential expression (*p*-value < 0.01) with Mtb *mtrA* knockdown of the conserved protein Rv1815 (log2 fold-change = -1.9) and the resuscitation promoting factors *rpfA* (log2 fold-change = -1.9), and *rpfC* (log2 fold-change = -2.5). Interestingly, the *rpfA* and *rpfC* protein products also have predicted PG cleavage activity^57,58^. Additionally, there was significant repression of *ripB* (*Rv1478*, log2 fold-change = -3.4), an PG endopeptidase with activity similar to *ripA*^*32*^, upon *mtrA* knockdown in Mtb. While these PG hydrolases are not part of corem 71 and may not be important for PG cleavage during infection in the host, they could be regulated by MtrA and active in other conditions. This points to another capability of EGRIN 2.0, to decipher how regulation by a TF varies conditionally across different targets.

Importantly, this study is the first to characterize MtrA loss directly in Mtb. Previous attempts to construct a *ΔmtrA* mutant in Mtb were unsuccessful^21,59,60^, but the CRISPRi platform for mycobacteria allowed easy manipulation of the essential gene^27^. Using CRISPRi to repress *mtrA* in Mtb and profile genome-wide transcriptional changes, we have validated new regulatory targets of the essential response regulator (**Table 1**). Moreover, we have conducted parallel experiments in Msm and found conserved MtrA regulatory targets between the species. However, there were also key differences in PG cleavage genes. (e.g., *Rv2525c, rpfB, rpfE*). These results, along with the important difference in MtrA essentiality between Msm and Mtb calls into question using the fast-growing mycobacterial model organism to interpret MtrA activity in Mtb. This caution may extend to the study of all mycobacterial PG remodeling enzymes, as there are fundamental differences in PG incorporation^61^ and PG hydrolase activity^52,62^ between Msm and Mtb. These findings propose evolution from saprophyte to pathogen has occurred, in part, through regulatory networks and PG remodeling activities to function in the host environment.

Using sgRNAs of various “strengths”, we were able to tune the magnitude of *mtrA* knockdown and establish that even minimal *mtrA* knockdown is lethal to Mtb. All bacteria need to coordinate synthesis and breakdown of PG to maintain integrity through cell division; the importance of this is evident from the number of PG-targeting antibiotics. However, the complexity of the Mtb cell wall represents a distinct challenge to the pathogen, requiring specialized mechanisms for cell division and growth to occur^63^. We propose the unique cell wall properties of Mtb and the regulation of multiple PG cleavage enzymes contributes to the extreme vulnerability of MtrA in Mtb. Furthermore, we demonstrated that <10% inhibition via *mtrA* knockdown, resulted in complete clearance of Mtb from cultures with sublethal doses of drugs currently used to treat TB. Ultimately, our work presents MtrA as an attractive therapeutic target with several advantages: (1) inhibition of MtrA potentiates frontline drugs, requiring lower therapeutic doses; (2) inhibition of MtrA restricts bacterial growth, especially within host environments permissive for varying degrees of growth; and (3) chemical space already exists^53^, making MtrA a tractable target. We hope that deciphering conditional gene regulatory networks in the Mtb EGRIN 2.0 model will identify additional drug targets and treatment-enhancing strategies to better control TB.

## Acknowledgements

We gratefully acknowledge Sarah Fortune for kindly providing us with the PLJR962 and PLJR965 plasmids. We thank Michael VanNieuwenhze for kindly providing HADA. We thank members of the Baliga lab for critical discussions and feedback. This work is funded by the National Institute of Allergy and Infectious Diseases of the National Institutes of Health (R01AI128215, R01AI141953 and U19AI135976) and the Bill and Melinda Gates Foundation (INV-009322).

## Author Contributions

EJRP and NSB conceived the study. ANB and DJR ran cMonkey and constructed the ensemble. EJRP analyzed the ensemble and all data represented. WW provided assistance with ensemble analyses and Web portal. RM analyzed Tn-seq data. ST developed the Web portal. EJRP designed and supervised experimental validations. AK, VS, WC, MP and RR performed experiments. MLAO helped with computational analyses related to rare co-regulation events. AB helped with designing experimental validations. EJRP and NSB wrote the paper and supervised the study. All authors read the manuscript and approved its content.

## Data availability

The data reported in this paper are available in Supplementary Datasets 1-5. EGRIN 2.0 code is available in Baliga lab GitHub (https://github.com/baliga-lab/egrin2-tools) with additional information on the supporting Web site (http://egrin2.systemsbiology.net). All corem and GREs of the Mtb EGRIN 2.0 model are available on the Mtb web portal: http://networks.systemsbiology.net/mtb/. The raw Tn-seq fastq sequence data files are deposited in the Sequence Read Archive database under accession PRJNA701946. The RNA-seq data generated for this study are available in the Gene Expression Omnibus under accession no. GSE166806.

## METHODS

### EGRIN 2.0 construction

EGRIN 2.0 was constructed as an ensemble of many individual EGRIN models (∼300 for *M. tuberculosis*). Each EGRIN model was constructed using *cMonkey2*^*4*^. The input to *cMonkey2* was 1,861 transcriptome profiles with metadata collected about each experiment to annotate the environmental context (termed condition block, **Dataset S5**). Other input to *cMonkey2* were upstream regions of all genes, and functional association networks, including operon predictions from MicrobesOnline^64^ and functional protein interactions from EMBL String databases^65^. We integrated the EGRIN models and mined the ensemble to discover frequently reoccurring features and associations as described previously^1^. We refer to the modules detected by our procedure as co-regulated modules, or corems, the frequently re-occurring de novo cis-regulatory motifs as gene regulatory elements, or GREs. Full description of the algorithms and each post-processing step is documented in Supplementary Information of Brooks *et al*^*1*^.

### Bacterial strains

All *M. tuberculosis* strains are derivatives of H37Rv; all *M. smegmatis* strains are derivatives of mc^2^155.

### Media

*M. tuberculosis* and *M smegmatis* were grown at 37°C in Middlebrook 7H9 broth or 7H10 plates supplemented with 0.2% glycerol, 0.05% Tween-80, and 10% ADC (liquid media) or OADC (plates), with aeration. Where indicated, anhydrotetracycline (ATc) was used at 100 ng/ml. For comparison of neutral and acidic pH media, the neutral media was 7H9 broth supplemented with 0.2% glycerol, 0.05% tyloxapol, and ADC buffered with 100 mM 3-(N-morpholino)propanesulfonic acid (MOPS) to pH 7. The acidic pH media was the same 7H9-rich media buffered with 100 mM 2-(N-morpholino)ethanesulfonic acid (MES) to pH 5.6. Frozen cells were inoculated into standard 7H9-rich media, grown to mid-log phase (OD_600_ ∼0.5-0.7), washed in 1x PBS three times, and diluted into either neutral or acidic pH media at a starting density of OD_600_ = 0.05.

### Transposon mutant library sequencing

Transposon mutant libraries were constructed using the ϕMycoMarT7 phagemid in *M. tuberculosis* H37Rv. The mutant libraries were grown to mid-log phase in 7H9-rich media, then diluted to OD_600_ = 0.1 in 7H9-rich media at neutral (pH 7.0) or acidic pH (pH 5.6) media. In another experiment, the mutant libraries were diluted back in 7H9-rich media with or without 0.05% SDS. Following dilution, mutant samples were diluted in PBS with 0.05% Tween-80 and plated onto 245 mm x 245 mm 7H10 plates supplemented with kanamycin (50 μg/ml) to T0. Additional samples were exposed to the stressors for 72 hours and then plated. Colonies grew for 21 days and were collected for genomic DNA isolation. The mutant composition was determined by sequencing amplicons of the transposon-genome junctions following the protocol outlined by Long *et al*^66^. Paired-end reads were run on an Illumina HiSeq 2500 at the Genomics Services Core at Fred Hutchinson Cancer Research Center. Mapping and quantification of transposon insertions sites was done using *TRANSIT* analysis platform^20^. The change in fitness between Day 3 and T0 for all genes was determined following the strategy of van Opijnen *et al*.^*67*^ using a custom processing pipeline, full description and code is available at http://networks.systemsbiology.net/mtb/. The raw Tn-seq fastq sequence data files are deposited in the Sequence Read Archive database under accession PRJNA701946.

### Permutation test for evaluating significance of overlap between corems and genes with reduced fitness

The genes with reduced delta fitness in stress (either acidic pH or SDS treatment) were permutated 1000 times to generate shuffled gene clusters. In each permutation, the produced shuffled gene clusters had the same size as corem 71. Then, the average delta fitness for each shuffled gene set was computed and compared to the average delta fitness for corem 71. The overall permutation test *p*-value was computed as the proportion of cases (out of 1000 permutations) in which the average delta fitness was equal or lower than the observed value in corem 71.

### Gene expression profiling of CRISPRi-mediated mtrA knockdown

For each sample, cultures were grown to mid-log phase in 7H9-rich media with Kanamycin (50 μg/ml) and then diluted back in the presence or absence of ATc to induce CRISPRi-mediated *mtrA* knockdown. Knockdown was allowed to proceed for 14 hours (*M. smegmatis*) or 4 days (*M. tuberculosis*). Samples, in biological quadruplicate, were collected by centrifugation at high speed for 5 min, discarding supernatant and immediately flash freezing the cell pellet in liquid nitrogen. Cell pellets were stored at -80° C until RNA extraction was performed as previously described^41^. The processing of samples for sequencing and read alignment is described below. The RNA-seq data profiling CRISPRi-mediated *mtrA* knockdown generated for this study are publicly available from the Gene Expression Omnibus at GSE166806.

### Gene expression profiling of mtrA overexpression

To investigate gene expression changes of *mtrA* overexpression, we used *M. tuberculosis* strain containing an ATc-inducible vector as described previously^17^. For each sample, cultures of *M. tuberculosis* were cultured in standard 7H9-rich media containing 50 μg/ml hygromycin B to mid-log phase, washed three times in 1x PBS and diluted back to OD_600_ = 0.05 into either neutral or acidic pH media with 50 μg/ml hygromycin B and the presence or absence of ATc inducer. Samples, in biological triplicate were collected after 18 hours of ATc induction, centrifuged at high speed for 5 min, supernatant was discarded and the cell pellet was immediately flash frozen in liquid nitrogen. Cell pellets were stored at -80° C until RNA extraction was performed as previously described^41^. The processing of samples for sequencing and read alignment is described below. The RNA-seq data profiling *mtrA* overexpression in netural and acidic pH generated for this study are publicly available from the Gene Expression Omnibus at GSE166806.

### Gene expression profiling at netural or acidic pH

To investigate gene expression changes of *M. tuberculosis* at acidic pH, cultures of *M. tuberculosis* were cultured in standard 7H9-rich media to mid-log phase, washed three times in 1x PBS and diluted back to OD_600_ = 0.1 into either neutral or acidic pH media. Samples, in biological triplicate, were collected right after transfer and after 24 hours in neutral or acidic pH media. Samples were centrifuged at high speed for 5 min, supernatant was discarded and the cell pellet was immediately flash frozen in liquid nitrogen. Cell pellets were stored at -80° C until RNA extraction was performed as previously described^41^. The processing of samples for sequencing and read alignment is described below. The RNA-seq data profiling *mtrA* overexpression in netural and acidic pH generated for this study are publicly available from the Gene Expression Omnibus at GSE166806.

### Processing and analysis of RNA-seq data

Sample collection and RNA-extraction was performed as described above. Total RNA samples were depleted of ribosomal RNA using the Ribo-Zero Bacteria rRNA Removal Kit (Illumina, San Diego, CA). Quality and purity of mRNA samples was determined with 2100 Bioanalyzer (Agilent, Santa Clara, CA). Samples were prepared with TrueSeq Stranded mRNA HT library preparation kit (Illumina, San Diego, CA). All samples were sequenced on the NextSeq sequencing instrument in a high output 150 v2 flow cell. Paired-end 75 bp reads were checked for technical artifacts using Illumina default quality filtering steps. Raw FASTQ read data were processed using the R package DuffyNGS^42^. Briefly, raw reads were passed through a 2-stage alignment pipeline: (i) a pre-alignment stage to filter out unwanted transcripts, such as rRNA; and (ii) a main genomic alignment stage against the genome of interest. Reads were aligned to *M. tuberculosis H37Rv* (ASM19595v2) or *M. smegmatis mc*^*2*^*155* (ASM1500v1) with Bowtie2^43^, using the command line option “very-sensitive.” BAM files from stage (ii) were converted into read depth wiggle tracks that recorded both uniquely mapped and multiply mapped reads to each of the forward and reverse strands of the genome(s) at single-nucleotide resolution. Gene transcript abundance was then measured by summing total reads landing inside annotated gene boundaries, expressed as both RPKM and raw read counts. All RNA-seq data (raw and processed data) generated for this study are publicly available at the Gene Expression Omnibus under accession numbers GSE166806.

### Differential expression analysis

We used a panel of 5 DE tools compiled in DuffyNGS to identify gene expression changes as previously described. The tools included (i) RoundRobin (in-house); (ii) RankProduct^68^; (iii) significance analysis of microarrays (SAM)^69^; (iv) EdgeR^70^; and (v) DESeq2^71^. Each DE tool was called with appropriate default parameters and operated on the same set of transcription results, using RPKM abundance units for RoundRobin, RankProduct, and SAM and raw read count abundance units for DESeq2 and EdgeR. Each DE tool’s explicit measurements of differential expression (fold-change) and significance (*p*-value) were similarly combined via appropriate averaging (arithmetic and geometric mean, respectively).

### Peptidoglycan labeling with fluorescent d-alanine analogues

Fluorescent d-alanine analog (FDAA) HADA (HCC-amino-d-alanine) was synthesized by Michael VanNieuwenhze at Indiana University using methods previously published^72^. Cultures were grown to mid-log phase in 7H9-rich media with Kanamycin (50 μg/ml) and then diluted back in the presence or absence of ATc to induce CRISPRi-mediated *mtrA* knockdown. Knockdown was allowed to proceed for 14 hours (*M. smegmatis*) or 4 days (*M. tuberculosis*). Cultures were then inoculated at an OD600 of 0.1 to 0.3 into 7H9 medium supplemented with 1 mM HADA and incubated for 3.5 hours (*M. smegmatis*) or 20 hours (*M. tuberculosis*). Bacterial suspension were washed 3 times with PBS-0.05% Tween-80 and fixed with paraformaldehyde for 30 min (*M. smegmatis*) or 4 hours (*M. tuberculosis*), to ensure bacterial death for further imaging outside a contained environment.

### Super-resolution microscopy

Fixed bacterial suspensions were mixed with the same volume of mounting medium and 10 μl amounts were spread on microscope slides and covered with cover glasses. Microscopy imaging was performed using SP8 Lightning super-resolution microscope (Leica Microsystems). Images were analyzed using ImageJ software.

### Phenotyping CRISPRi-mediated mtrA knockdown

To visually monitor the effects of CRIPSRI-mediated mtrA knockdown, cultures were grown to mid-log phase in 7H9-rich media with Kanamycin (50 μg/ml) and spotted on solid media with or without Atc. Plates were incubated at 37°C for 21 days (*M. tuberculosis*).

### Susceptibility to antibiotics assay

The CRISPRi-mediated *mtrA* knockdown strain and control sgRNA of *M. smegmatis* was spread on LB agar plates containing 100 ng/ml anhydrotetracycline inducer. A filter disc with 10 μl of carbenicillin (100 mg/ml), isoniazid (0.5 mg/ml), vancomycin (6 mg/ml), cycloserine (100 mg/ml) or rifampicin (0.5 mg/ml) was placed in the center of plate and the diameter of inhibition of growth was measured after 4 days of growth.

### Time-kill assay

For each sample, *M. tuberculosis* cultures were grown to mid-log phase in 7H9-rich media with Kanamycin (50 μg/ml) and then diluted back in the presence or absence of ATc to induce CRISPRi-mediated *mtrA* knockdown. Knockdown was allowed to proceed for 4 days. Antibiotic (0.18 μg/ml isoniazid or 0.006 μg/ml) or vehicle control was added to the cultures and samples were serially diluted and plated on 7H10 agar plates. Samples were taken for 7 days following antibiotic treatment and colonies were counted after 21 days. All time-kill assays were performed in biological triplicate.

