## Supplemental Material for "MtrA regulation of essential peptidoglycan cleavage in *Mycobacterium tuberculosis* during infection"

### List of Figures

**Figure S1.** Corem 71 fitness in acidic pH and cell wall-perturbing detergent SDS.

**Figure S2**. Cell viability of CRISPRi knockdown of *mtrA* with “strong” PAMs in *M. tuberculosis.*

**Figure S3**. Cell viability of CRISPRi knockdown of *mtrA* with PAMs of various “strengths” in *M. tuberculosis*.

**Figure S4**. Growth of CRISPRi knockdown of *mtrA* with “strong” PAMs in *M. smegmatis.*

**Figure S5.** Volcano plot of differentially expressed genes for induced vs uninduced CRISPRi knockdown of *mtrA* in *M. smegmatis*.

**Figure S6.** MtrA controls cell division in *M. smegmatis*.

### List of Tables

**Table S1.** sgRNAs used in this study.

**Table S2.** MtrA regulatory targets identified across ChIP-seq studies.

**List of Datasets (separate files)**

**Dataset S1.** Delta fitness in stress conditions (acidic pH and SDS treatment) for all genes from Tn-seq analysis.

**Dataset S2.** Results of overlap between corems and genes with reduced fitness in acidic pH (and results of same analysis with SDS treatment).

**Dataset S3.** Results of differential expression results of CRISPRi knockdown of *mtrA* in *M. tuberculosis*.

**Dataset S4.** Results of differential expression results of CRISPRi knockdown of *mtrA* in *M. smegmatis*.

**Dataset S5.** Compendia of gene expression data used as input for EGRIN 2.0 and their annotated condition block.

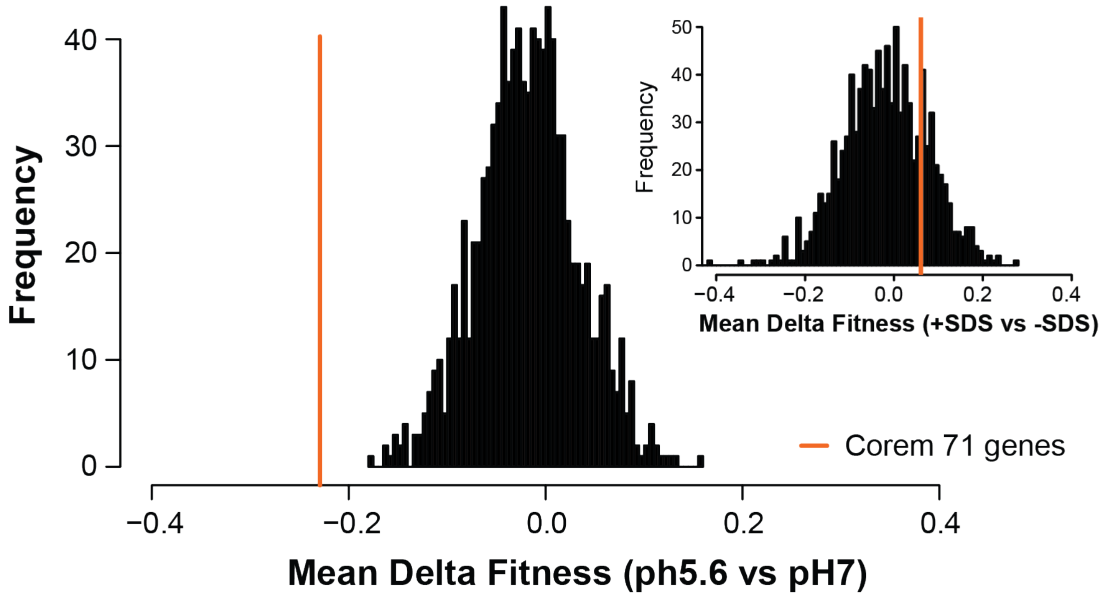


**Figure S1. Corem 71 fitness in acidic pH and cell wall-perturbing detergent SDS.** Histogram of mean delta fitness between acidic pH and neutral pH from 1000 permutations to generate shuffled gene sets. In each permutation, the produced shuffled gene set had the same size as corem 71. The orange line represents the observed value for corem 71. Inset displays the results of same analysis with mean delta fitness between the presence and absence of 0.05% SDS.

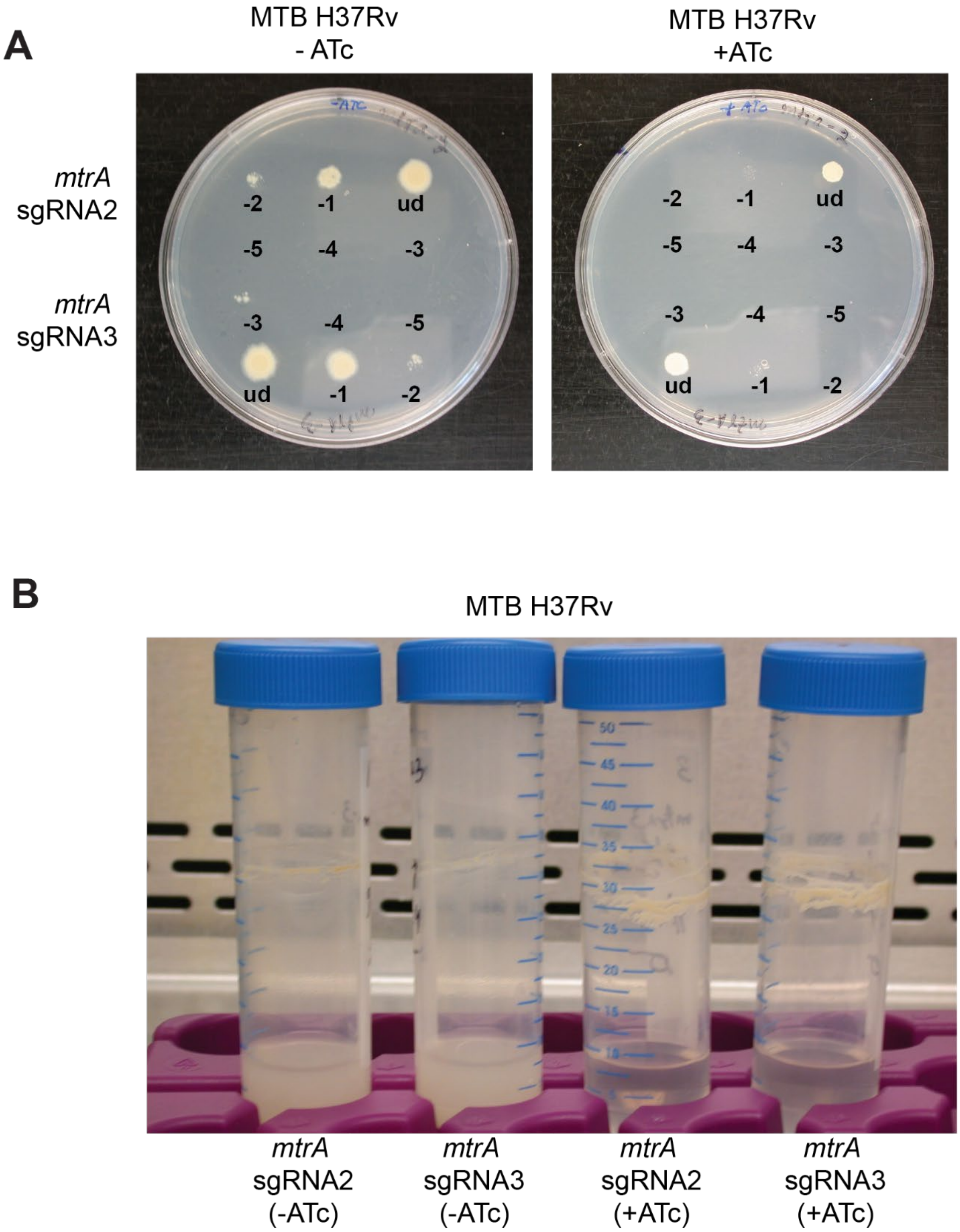


**Figure S2. Cell viability of CRISPRi knockdown of *mtrA* with "strong" PAMs in *M. tuberculosis*.** (**A**) Serial 10-fold dilutions of *M. tuberculosis* H37Rv CRISPRi strains with sgRNA2 and sgRNA3 were spotted on 7H10 agar plates with or without ATc. (**B**) Photographs of liquid cultures of the indicated strains.

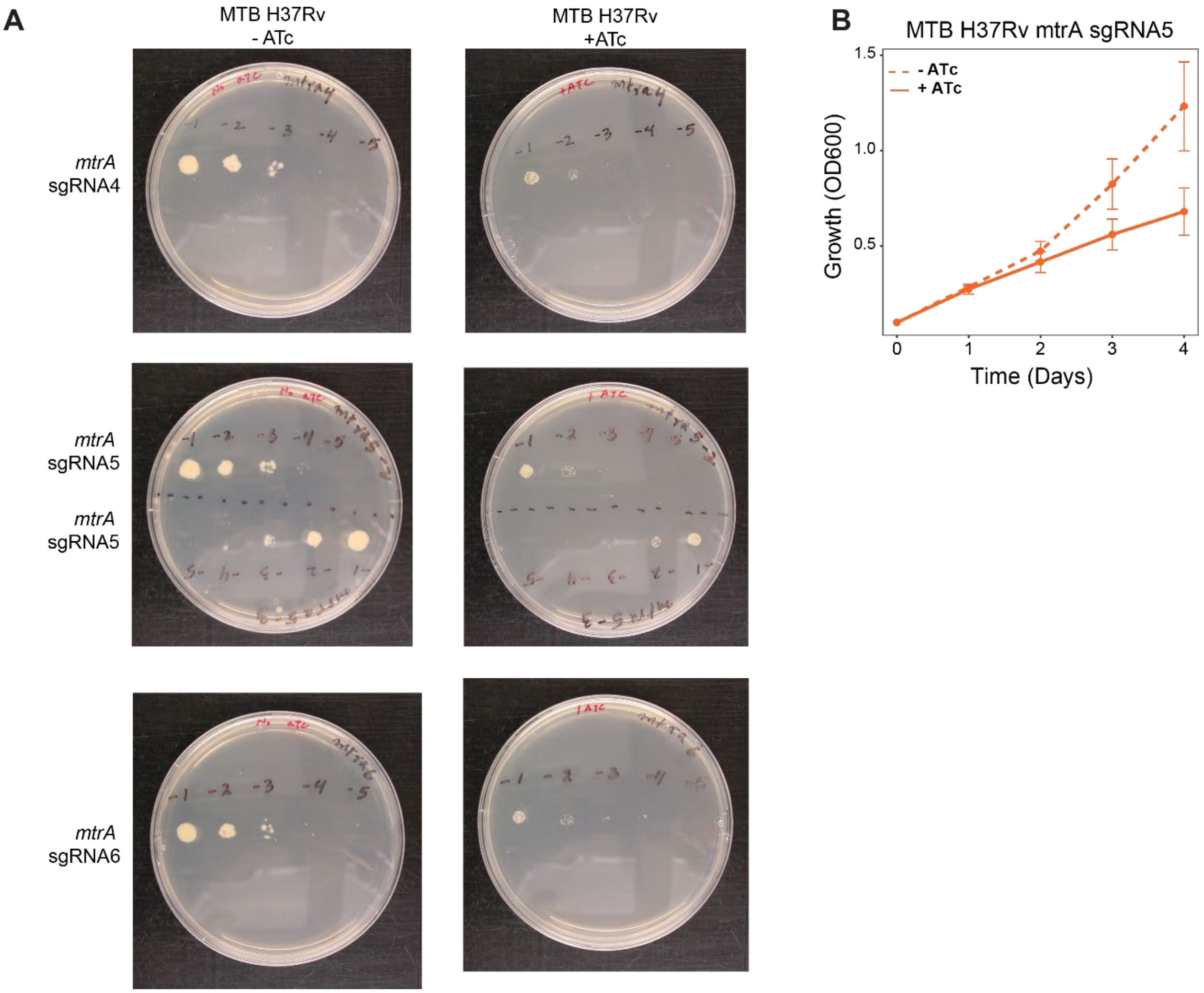


**Figure S3. Cell viability of CRISPRi knockdown of *mtrA* with PAMs of various "strengths" in *M. tuberculosis*.** (**A**) Serial 10-fold dilutions of *M. tuberculosis* H37Rv CRISPRi strains with sgRNA4, sgRNA5 and sgRNA6 were spotted on 7H10 agar plates with or without ATc. (**B**) Growth of *M. tuberculosis* H37Rv CRISPRi strain with sgRNA5 in liquid 7H9-rich media with (solid line) or without (dotted line) ATc. Growth was monitored daily by optical density at 650 nm. Points are the average of three biological replicates and error bars represent standard deviation.

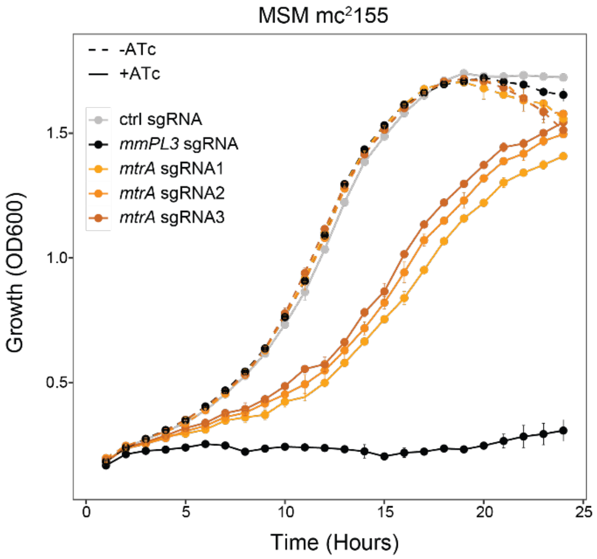


**Figure S4. Growth of CRISPRi knockdown of *mtrA* with "strong" PAMs in *M. smegmatis.*** Growth of *M. smegmatis* $mc^2155$ CRISPRi strains with sgRNA1, sgRNA2, sgRNA3, control (ctrl) sgRNA, and sgRNA targeting the essential gene *mmPL3* in liquid 7H9-rich media with (solid line) or without (dotted line) ATc. Growth was monitored hourly by optical density at 650 nm. Points are the average of three biological replicates and error bars represent standard deviation.

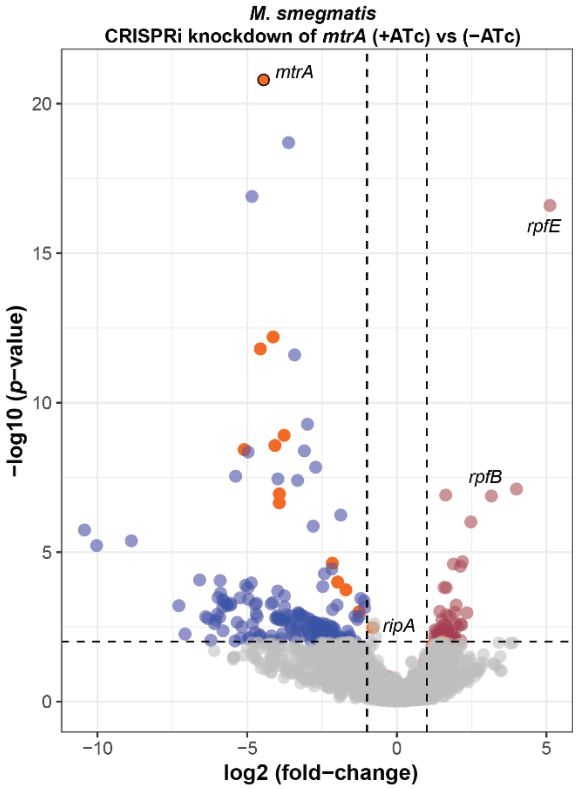


**Figure S5. Volcano plot of differentially expressed genes for induced vs uninduced CRISPRi knockdown of *mtrA* in *M. smegmatis*.** The significantly differentially expressed genes were selected by $p$-value < 0.01 and absolute log2 fold-change > 1. Dots represent different genes, with labels for particular genes of interest. Grey dots are genes without significant different expression, red dots are significantly up-regulated genes ($N$ = 58 genes) and blue dots are significantly down-regulated genes ($N$ = 185 genes). The orange dots are all genes of corem 71.

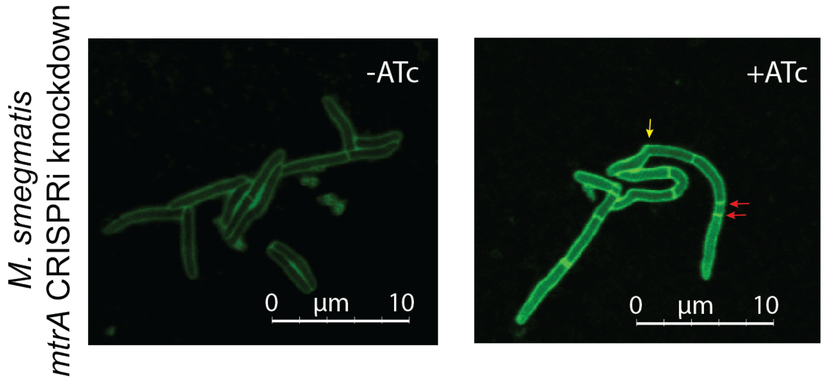


**Figure S6. MtrA controls cell division in *M. smegmatis*.** Example micrographs of uninduced (-ATc) and induced (+ATc) CRISPRi knockdown of *mtrA* with sgRNA1 in Msm. After knockdown, cells were labeled with HCC-amino-D-alanine (HADA) for 3.5 h. Red arrows indicate multiple septa and yellow arrow indicates the curved shape phenotype. Data are representative of at least two independent experiments.

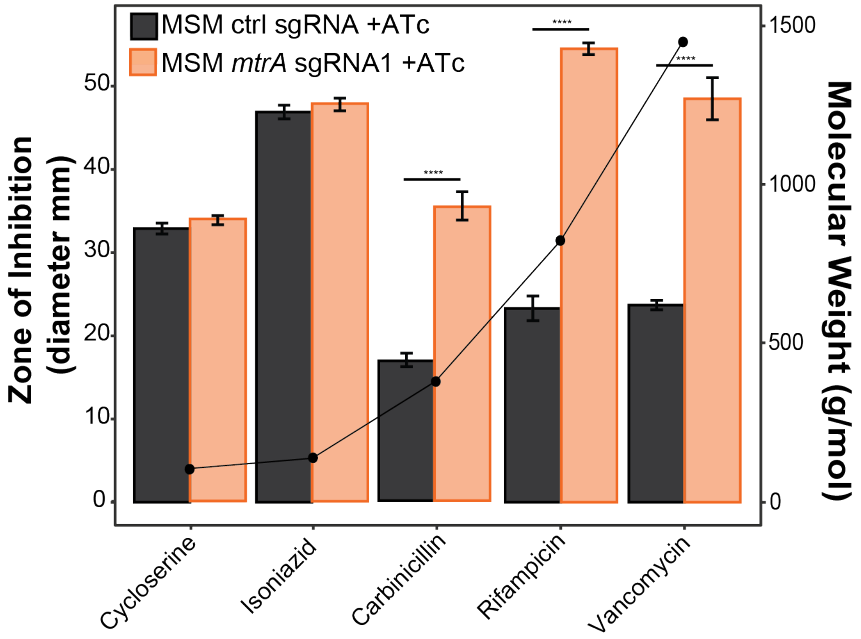


**Figure S7. The knockdown of *mtrA* increases sensitivity to high molecular weight antibiotics in *M. smegmatis*.** The induced CRISPRi *mtrA* knockdown with sgRNA1 and control (ctrl) sgRNA strains of Msm were spread on LB plates containing 100 ng/ml ATc inducer in triplicate. A filter disc with 10 µl antibiotic was place in the center of plate and the diameter of inhibition of growth was measured after 4 days of growth. Antibiotic concentration on disc: cycloserine (100 mg/ml); isoniazid (X); carbinicillin (100 mg/ml); rifampicin (X); vancomycin (6 mg/ml). Significance was determined by Student's T-test. ****: $p$-value < 0.0001. Data are representative of at least two independent experiments.

**Table S2**. **sgRNAs used in this study**.

| Organism | Gene | sgRNA name | Forward | Reverse | PAM (5'-3') | Fold-repression |
|---|---|---|---|---|---|---|
| *M. smegmatis* | *mtrA* | sgRNA1 | GGGAGGTGTCGAAACCCTCACCAC | AAACGTGGTGAGGGTTTCGACACC | GTTCT | 120.5 |
| *M. smegmatis* | *mtrA* | sgRNA2 | GGGAATCGATGCCGTTCATCCCAG | AAACCTGGGATGAACGGCATCGAT | ATGCT | 84.6 |
| *M. smegmatis* | *mtrA* | sgRNA3 | GGGAGACGGTGTCGGTCTTGGCGG | AAACCCGCCAAGACCGACACCGTC | ATGCT | 84.6 |
| *M. smegmatis* | *n/a* | control_sgRNA | GGGAGGAGACGATTAATGCGTCTCG | AAACCGAGACGCATTAATCGTCTCC | TTTCT | 158.1 |
| *M. smegmatis* | *mmpL3* | mmpL3_sgRNA | GGGAGCGACAGACTGGCTGCCCTCGTC | AAACGACGAGGGCAGCCAGTCTGTCGC | TTTCT | 158.1 |
| *M. tuberculosis* | *mtrA* | sgRNA2 | GGGAATCCACGGTGTCGGTCTTTGCGG | AAACCCGCAAAGACCGACACCGTGGAT | ATGCT | 84.6 |
| *M. tuberculosis* | *mtrA* | sgRNA3 | GGGAGATTTCTACGTCGGCGATGG | AAACCCATCGCCGACGTAGAAATC | ATGCT | 84.6 |
| *M. tuberculosis* | *mtrA* | sgRNA4 | GGGAGTCTTTGCGGTGAGCATCACGA | AAACTCGTGATGCTCACCGCAAAGAC | GTTCC | 51.5 |
| *M. tuberculosis* | *mtrA* | sgRNA5 | GGGAGCCGGTAACCCCATACCTGTT | AAACAACAGGTATGGGGTTACCGGC | CTGCT | 42.2 |
| *M. tuberculosis* | *mtrA* | sgRNA6 | GGGAGTCAGCACCACAGTCGGGTTC | AAACGAACCCGACTGTGGTGCTGAC | ATCCC | 24.2 |

**Table S2. MtrA regulatory targets identified across ChIP-seq studies.** Summary of the 14 putative MtrA targets identified across studies that have evaluated MtrA regulatory targets using global ChIP-seq analysis (Minch et al., 2015; Chatterjee et al., 2018; Gorla et al., 2018) and gene expression from CRISPRi *mtrA* knockown in Mtb.

| Gene | Name | Description | Mtb *mtrA* knockdown +ATc vs -ATc Log2 FC | Mtb *mtrA* knockdown +ATc vs -ATc *p*-value | Mtb *mtrA* knockdown strain |
|---|---|---|---|---|---|
| *Rv1884c* | *rpfC* | resuscitation-promoting factor RpfC | -2.87262 | 1.02E-14 | sgRNA2 |
| *Rv1815* | | hypothetical protein | -2.32863 | 4.96E-05 | sgRNA2 |
| *Rv1884c* | *rpfC* | resuscitation-promoting factor RpfC | -2.1646 | 1.14E-56 | sgRNA3 |
| *Rv0867c* | *rpfA* | resuscitation-promoting factor RpfA | -1.87012 | 2.76E-23 | sgRNA3 |
| *Rv0867c* | *rpfA* | resuscitation-promoting factor RpfA | -1.85562 | 0.000101 | sgRNA2 |
| *Rv1815* | | hypothetical protein | -1.50218 | 1.26E-27 | sgRNA3 |
| *Rv1886c* | *fbpB* | mycolyltransferase Ag85B | -0.33581 | 0.000702 | sgRNA2 |
| *Rv1918c* | *PPE35* | PPE family protein PPE35 | 0.736914 | 3.39E-06 | sgRNA3 |
| *Rv1116* | | hypothetical protein | 3.48704 | ns | sgRNA2 |
| *Rv1116* | | hypothetical protein | 1.602459 | ns | sgRNA3 |
| *Rv1918c* | *PPE35* | PPE family protein PPE35 | 0.765059 | ns | sgRNA2 |
| *Rv1268c* | | hypothetical protein | 0.684585 | ns | sgRNA3 |
| *Rv1886c* | *fbpB* | mycolyltransferase Ag85B | -1.04448 | ns | sgRNA3 |
| *Rv3857c* | | Membrane protein | -0.25223 | ns | sgRNA2 |
| *Rv1009* | *rpfB* | resuscitation-promoting factor RpfB | -0.70509 | ns | sgRNA2 |
| *Rv1268c* | | hypothetical protein | 0.921308 | ns | sgRNA2 |
| *Rv3857c* | | membrane protein | -0.98162 | ns | sgRNA2 |
| *Rv1361c* | *PPE19* | PPE family protein PPE19 | -0.21558 | ns | sgRNA3 |
| *Rv1523* | | methyltransferase | -0.58467 | ns | sgRNA2 |
| *Rv0116c* | *ldtA* | L,D-transpeptidase LdtA | -0.09104 | ns | sgRNA2 |
| *Rv0950c* | | hypothetical protein | -0.29929 | ns | sgRNA2 |
| *Rv0950c* | | hypothetical protein | 0.015364 | ns | sgRNA3 |
| *Rv1361c* | *PPE19* | PPE family protein PPE19 | -0.12696 | ns | sgRNA2 |
| *Rv1523* | | methyltransferase | 0.103242 | ns | sgRNA3 |
| *Rv0116c* | *ldtA* | L,D-transpeptidase LdtA | 0.064224 | ns | sgRNA3 |
| *Rv1009* | *rpfB* | resuscitation-promoting factor RpfB | 0.26947 | ns | sgRNA3 |
| *Rv2352c* | *PPE38* | PPE family protein PPE38 | -0.25328 | ns | sgRNA2 |
| *Rv2352c* | *PPE38* | PPE family protein PPE38 | 0.059488 | ns | sgRNA3 |
